# One and Done: A safe, adaptable single-cycle SARS-CoV-2 vaccine platform blocks XBB.1.5 infection and transmission

**DOI:** 10.64898/2026.03.09.709481

**Authors:** Jacob Schön, Nico Joel Halwe, Tobias Britzke, Angele Breithaupt, Lorenz Ulrich, Jana Kochmann, Björn Corleis, Enja Tatjana Kipfer, Thomas Klimkait, Donata Hoffmann, Fabian Otte, Martin Beer, David Hauser

## Abstract

A next-generation SARS-CoV-2 vaccine must address the currently inadequate prevention of virus transmission, particularly against emerging variants of concern, a challenge that none of the licensed commercial vaccines fully meet. Effective control of respiratory pandemics necessitates vaccines that 1) can be rapidly adapted, 2) have high patient compliance with simple and non-invasive administration, and 3) block transmission in a virus challenge.

We describe here the characterization of an updated single-cycle SARS-CoV-2 vaccine candidate (scVac), engineered as a replication-defective virus with targeted deletions of the E gene and ORF6 and ORF7a, along with a truncation of ORF3a. The candidate carries an Omicron XBB.1.5 Spike (scVac^XBB^), maintaining all essential antigenic properties. The vaccine demonstrated an excellent safety profile in K18-hACE2 transgenic mice, the most sensitive virulence model, with no clinical signs or adverse events observed. In the Syrian hamster model, potent systemic and mucosal immune responses were induced, along with a strong neutralizing antibody response. Notably, there was no virus transmission to co-housed naïve animals, which outperforms a bivalent Omicron mRNA vaccine reference.

Our results demonstrate that scVac^XBB^-induced immunity not only prevents disease but also effectively blocks transmission. Furthermore, the successful introduction of the XBB.1.5 Spike protein into the scVac platform demonstrates the pipeline’s ability to adapt quickly to any emerging variant. These findings highlight the potential of this single-cycle concept as a next-generation COVID-19 vaccine, offering robust protection with a strong safety profile.

## Introduction

The unexpected global emergence of the severe acute respiratory syndrome coronavirus 2 (SARS-CoV-2) has underscored the general urgent need for innovative vaccine platforms capable of eliciting substantial and durable immune responses, especially against respiratory viral pathogens. Inactivated and mRNA-based vaccines have been pivotal in controlling the SARS-CoV-2 pandemic and in preventing severe disease outcomes. Nevertheless, they do not induce sufficient nasal mucosal immunity to block shedding or infection [1]. Historically, live-attenuated vaccines (LAVs) have offered a unique potential to induce comprehensive immune responses, encompassing both humoral and cellular immunity [2]. LAVs have provided robust and long-lasting immunity by mimicking natural infection without causing disease, making them a compelling option for vaccine development [3].

Among live vaccine approaches, single-cycle constructs, which are propagation-defective but antigenically complete, represent a promising option for safe and effective immunization [4–6]. A particularly innovative strategy in this realm involves the targeted deletion of essential genes, rendering the virus replication-defective after one initial round of infection. For coronaviruses, the Envelope (E) protein plays a crucial role in viral assembly and budding [7,8]. Deletion of the E gene thus severely impairs particle assembly and release, resulting in a virus capable of only a single round of infection. Such E-deletion constructs preserve the immunogenic properties of the virus while minimizing safety concerns, as they cannot produce infectious progeny virions [9–12].

In a previous study, we applied this strategy to SARS-CoV-2 and developed a single-cycle SARS-CoV-2 vaccine (scVac) candidate using an E-deletion construct, with additional deletions in immunologically relevant open reading frames (ORFs) [13]. In a proof-of-principle experiment, these constructs elicited high levels of protection and transmission-blocking immunity in preclinical models, validating their potential as a safe and efficacious vaccine candidate [13]. While it is widely assumed that live vaccines exhibit broad cross-protection, the ability to adjust, especially the spike gene, to the most recent circulating variants appears critical to maintain optimal protection against circulating SARS-CoV-2 variants.

Here, we report the development and characterization of a new SARS-CoV-2 scVac-construct that carries an updated XBB.1.5 Spike. We highlight its ideal safety properties and its potential as an adaptable platform technology. We studied the replication dynamics *in vitro* and its immunogenic potential *in vivo*.

## Results

### Vaccine generation and *in vitro* confirmation of single-cycle properties

Based on the previously described E-deleted single-cycle SARS-CoV-2 constructs and utilizing the CLEVER reverse genetics method [14], we generated a single-cycle vaccine (scVac) with deletions of the E, ORF6, and ORF7a genes. The additional insertion of a stop codon in ORF3a ensured viral genetic stability. The gene for the ancestral Spike protein was replaced with the counterpart of Omicron XBB.1.5. This construct was designated “scVac^XBB^“ (**Fig. 1a**). The genetic stability of the intended precise deletions in scVac^XBB^ was verified by repeated passaging on Vero-E2T cells (trans-complementing Vero E6 cell line that expresses E, ORF6, ORF7, and ORF8 [13], essential for scVac amplification) by both Next-Generation Sequencing (NGS) and Sanger sequencing. Its single-cycle properties were confirmed by sequential passages on several SARS-CoV-2 susceptible cell lines, Vero E6, Caco2, and A549-ACE2/TMPRSS2 cells side-by-side with a replication-competent wild-type SARS-CoV-2 carrying the XBB.1.5 spike (rCoV2^XBB^) (**Fig. 1b**).

**Figure 1:**
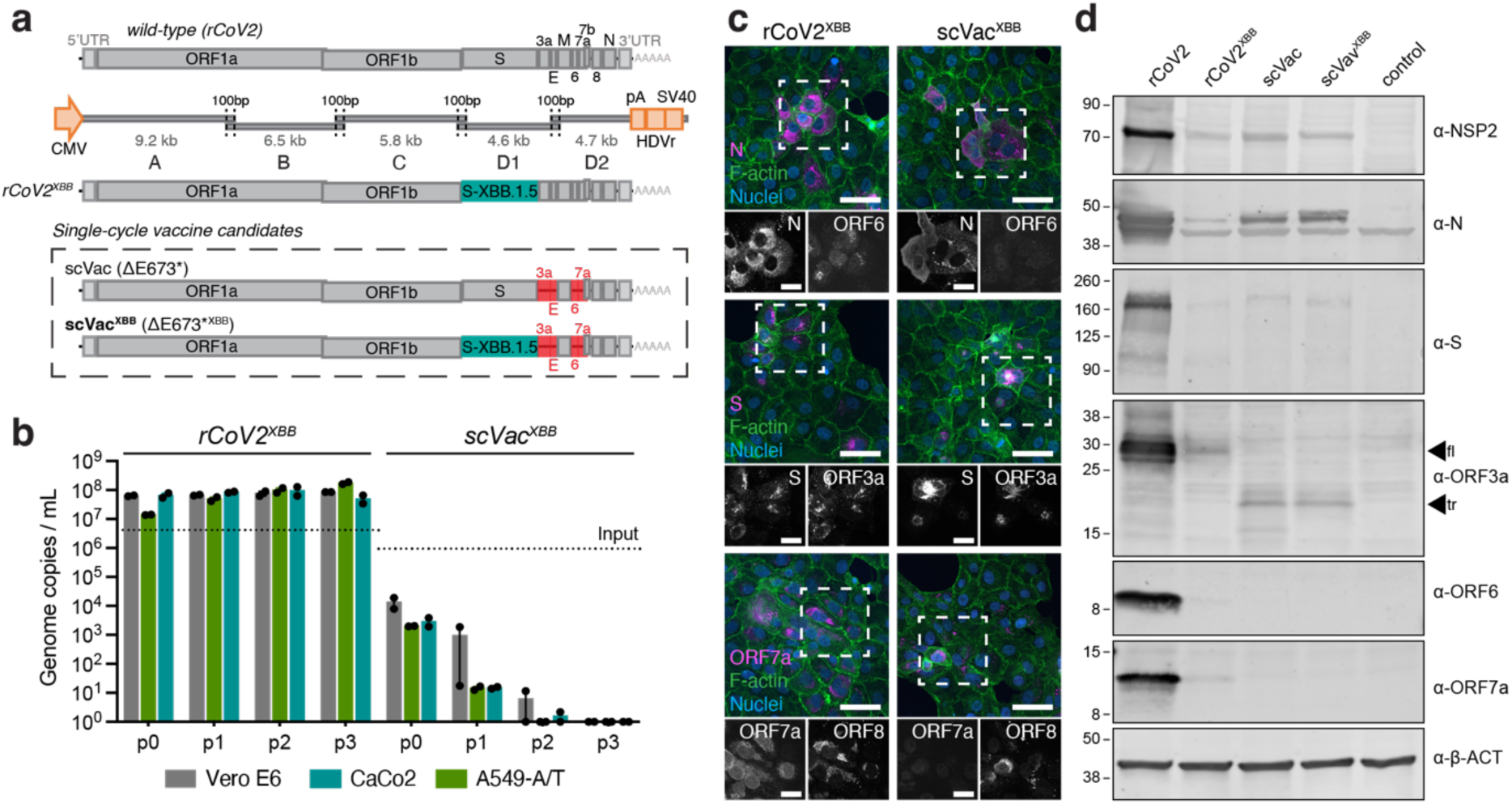
Genomic organization of the single-cycle vaccine, viral rescue, and *in vitro* characterization. (a) Schematic illustration of the SARS-CoV-2 genomic landscape and the deletions/substitutions in ΔE673*^XBB^ (scVac^XBB^); position of main structural and accessory proteins as indicated. Positions of the overlapping PCR fragments (A-D2) covering the whole SARS-CoV-2 genome are indicated. (b) Cell-free virus after passage of either wild-type SARS-CoV-2 expressing an Omicron XBB.1.5 Spike (rCoV2^XBB^) or scVac^XBB^ after 0-3 virus passages is shown for three non-complementing cell lines as indicated. The level of input virus is indicated by a dotted line. (c) Staining of N, S, and ORF7a (magenta), F-actin (green), nuclei (blue), and ORF6, ORF3a, or ORF8 in Vero E6-TMPRSS2 cells infected with rCoV2^XBB^ and scVac^XBB^. (d) Immunoblot analysis of viral proteins 24h after infection of Vero E6-TMPRSS2 cells with the indicated viruses or medium only (control), probed with anti-NSP2, anti-N, anti-S, anti-ORF3a (full-length [fl] and truncated [tr] forms indicated with arrows), anti-ORF6, anti-ORF7a, and anti-beta-actin (β-ACT) antibodies. Scale bar is 50 µm and 20 µm in (c) (overview and ROI images, respectively).

For further validation, the viral protein expression pattern of scVac^XBB^ was analyzed in infected Vero E6-TMPRSS2 cells (Vero E6 cells stably expressing TMPRSS2 [13]) by immunocytochemistry (**Fig. 1c**) and immunoblotting (**Fig. 1d**).

### Safety of vaccine candidate scVac^XBB^ and virulence analysis in the K18-hACE2 mouse model

To confirm the single-cycle properties and safety of scVac^XBB^, K18-hACE2 mice (n=16) were intranasally inoculated with 3x10^4^ infectious units (**Supplementary Fig. 1**). Survival, body weight, and viral genome levels were quantified in naso-buccal swabs and organ samples 2, 5, and 21 days post-inoculation (dpi) (**Fig. 2a**). Results were compared to a mock-inoculated control group and a SARS-CoV-2 Omicron XBB.1.5-infected group (10^3.73^ TCID_50_). No mortality and no body weight loss were observed in the scVac^XBB^- or in the mock-inoculated group, whereas mortality (two animals met human endpoint criteria at 7 and 10 dpi, counted as dead one day later in the survival curve) (**Fig. 2b**) and significant weight loss (comparison at 7 dpi, mock vs. SARS-CoV-2 XBB.1.5 [*p < 0.0001*]; scVac^XBB^ vs. SARS-CoV-2 XBB.1.5 [*p = 0.0002*]) occurred in the XBB.1.5-infected group (**Fig. 2c**). Viral genome quantification from combined naso-buccal swabs confirmed virus shedding and replication in the XBB.1.5-, but not in the scVac^XBB^ inoculated animals (similar to mock-inoculated groups) (**Fig. 2d**). In accordance, SARS-CoV-2-specific antibodies were detected only in the wild-type XBB.1.5-infected group (**Fig. 2e**). Lung tissue (8/12) and CNS samples (6/12) of XBB.1.5-infected animals contained high viral RNA yields (**Fig. 2f–i**), while in the scVac^XBB^-inoculated animals, only a low level of viral genomes was detectable in lung tissue (5/10) on days 2 and 5 after infection (**Fig. 2f and g**). No signal was detected in central nervous system (CNS) tissue, including cerebrum and cerebellum (0/10) (**Fig. 2h and i**).

**Figure 2:**
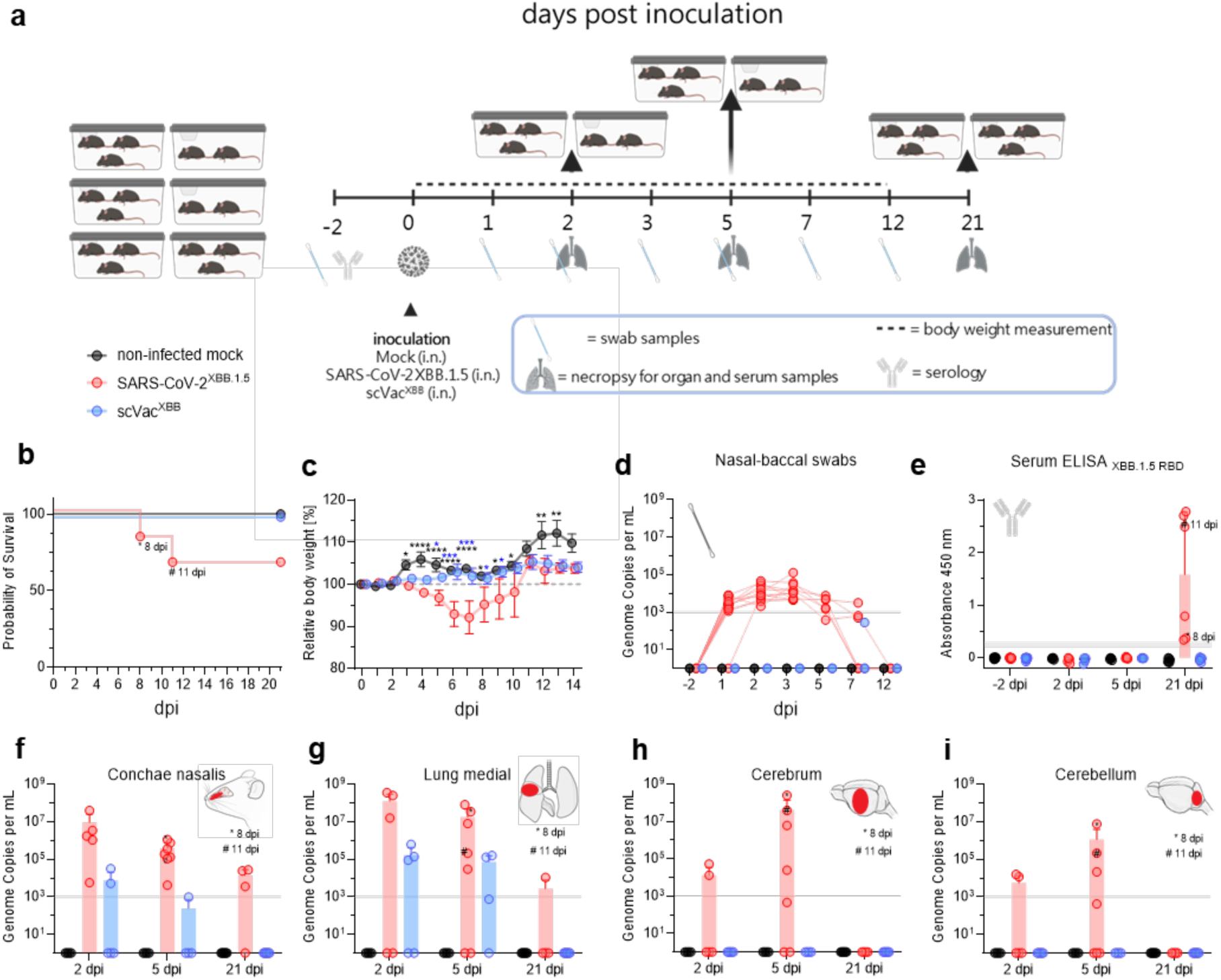
Intranasal inoculation of transgenic K18-hACE2 mice demonstrated an excellent safety profile of scVac^XBB^. (a) Experimental setup for sixteen K18-hACE2 mice per group after intranasal inoculation using the conditions indicated in grey (mock), red (wild-type), or blue (scVac). (b) Kaplan-Meyer curves of the survival in each vaccination group. (c) Relative body weight after inoculation. (d) Quantification of SARS-CoV-2 genome copies (ORF1b) in naso-buccal swabs (individual values). (e) XBB.1.5 Spike-specific RBD ELISA for samples taken at the indicated times after infection. (f-i) Quantification of SARS-CoV-2 genome copies (ORF1b) in (f) Concha nasalis, (g) Lung, (h) Cerebrum, and (i) Cerebellum tissue samples at 2, 5 (including the 8 and 11 dpi human endpoint animals), and 21 dpi. A grey line indicates quantification limits. Statistical differences between infected and non-infected groups were calculated using 2-way ANOVA with Tukey’s multiple comparison test. *\*p < 0.05, **p < 0.01, ***p < 0.001, ****p < 0.0001*. The mean is indicated for each group, with bars representing the SEM.

### scVac^XBB^: Transmission and immunogenicity in Syrian hamsters

To evaluate transmission and immunogenicity in comparison to an mRNA vaccine, Syrian hamsters (n=16 per group) were intranasally (i.n.) inoculated twice (prime-boost regimen) with either scVac^XBB^, mock (cell culture medium) or intramuscularly (i.m.) with an Omicron-based bivalent mRNA vaccine (BA.4/BA.5) (**Fig. 3a**). Naïve direct contact animals were co-housed in a 1:1 setting with 24-hour separation after prime and boost-vaccination to prevent nonspecific transmission of the inoculum (**Fig. 3a**). No mortality and no body weight loss were observed in any of the groups after prime or boost-vaccination (**Fig. 3b, c**) or in the respective naïve contact animals following co-housing (**Fig. 3d, e**).

**Figure 3:**
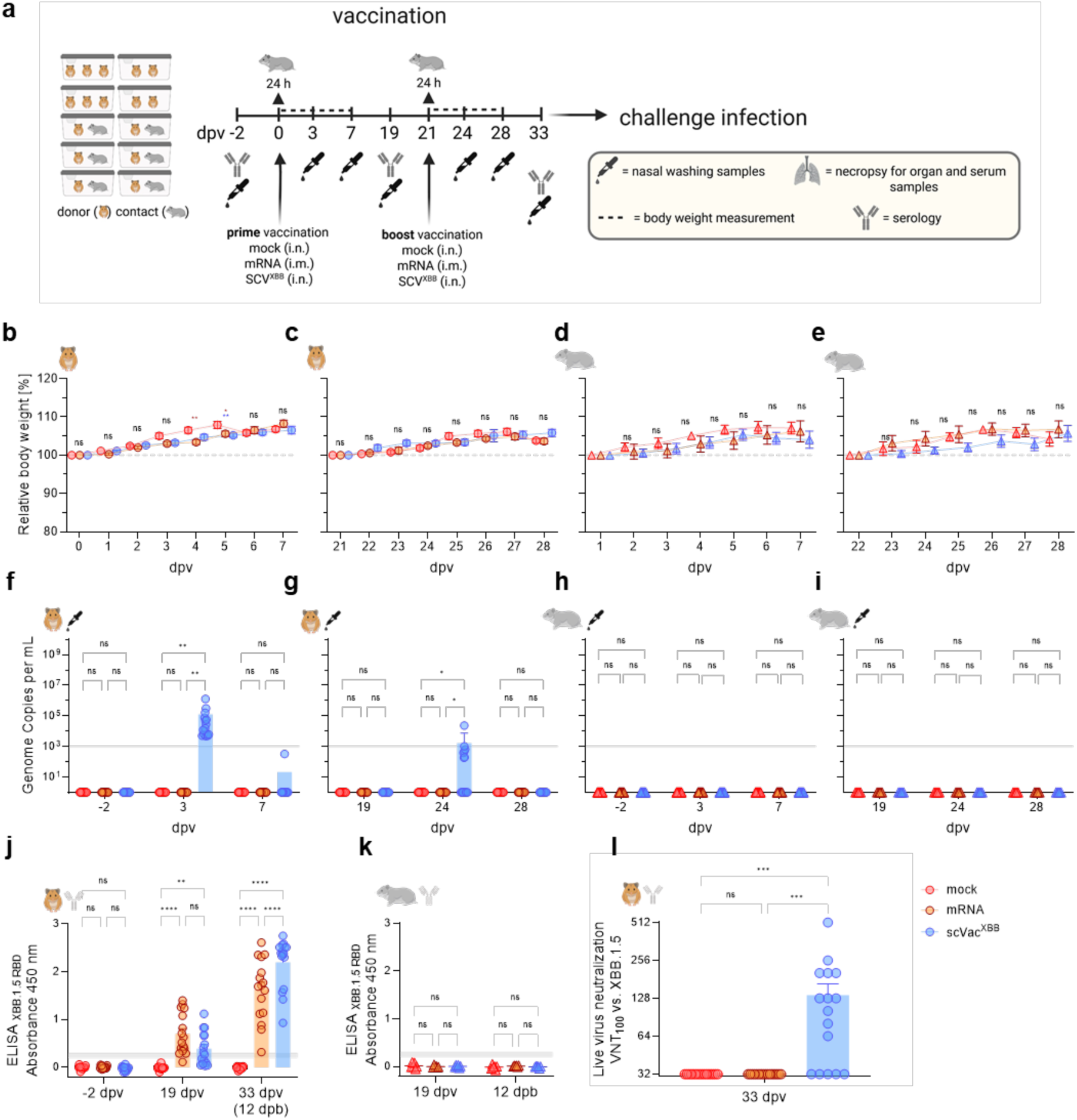
Intranasal vaccination with scVac^XBB^ confirms safety, including blocking transmission to direct contact hamsters. (a) Vaccination scheme of the prime-boost-vaccinated 16 hamsters that received either scVac^XBB^, a bivalent mRNA vaccine (BA.4/BA.5), or non-infectious cell culture medium (mock vaccination). Direct contact animals were separated for 24 hours at the time of vaccination in a 1:1 setup. (b-e) Body weight after prime or boost vaccination in vaccinated donor animals (b, c), and in naïve direct contact animals (d, e). (f-i) SARS-CoV-2 genome copies (ORF1b) per mL in nasal washing samples of vaccinated animals (donor) following prime (f) and boost (g) or of direct contact animals following prime (h) or boost (i) vaccination. (j, k) Humoral SARS-CoV-2 specific immune responses in sera of mRNA and scVac^XBB^-vaccinated animals (j), or naïve direct contact animals following co-housing (k). (l) SARS-CoV-2 neutralizing capacity against SARS-CoV-2 Omicron XBB.1.5 in sera of scVac^XBB^-, mRNA-, or mock-vaccinated animals before challenge infection. Statistical significance was calculated by repeated measures or by ordinary two-way ANOVA with Tukey’s or Bonferroni’s multiple comparison test. *\*p < 0.05, **p < 0.01, ***p < 0.001, ****p < 0.0001*, ns = non-significant. Besides individual values, group mean values are indicated by bars together with the standard error. The dotted gray line in b-e indicates starting body weight, and the gray line in f-k indicates threshold.

Quantification of genome copies in nasal washing samples revealed positive results for all scVac^XBB^-vaccinated animals at 3 days post-vaccination (dpv) (**Fig. 3f**) and only for one animal at 3 days post-boost (3 dpb, 24 dpv) (**Fig. 3g**) but none at 7 dpv (**Fig. 3f**) or 7 dpb (28 dpv) (**Fig. 3g**). None of the contact animals tested positive for viral genomes after co-housing post-prime (**Fig. 3h**) or post-boost (**Fig. 3i**). This indicates the absence of transmission of the single-cycle vaccine. A SARS-CoV-2 XBB.1.5-matching humoral immune response was detected as early as 19 dpv in both scVac^XBB^ and mRNA-vaccinated animals, with enhanced reactivity observed 12 dpb (**Fig. 3j**). The absence of scVac^XBB^ transmission was supported by the seronegativity of direct contact animals after both prime and boost vaccination (**Fig. 3k**). SARS-CoV-2 Omicron XBB.1.5 neutralizing activity was detected in 11 out of 16 scVac^XBB^-vaccinated animals at 12 dpb (33 dpv), but in none of the animals in the mRNA or mock-vaccinated groups (**Fig. 3l**).

Overall, these data further confirm the excellent safety of scVac^XBB^, with only a short, transient period of vaccine virus detectability and no transmission. Additionally, a robust humoral and neutralizing SARS-CoV-2-specific immune response was induced following vaccination.

### Protection and transmission of XBB.1.5 in challenge infections

Vaccinated hamsters were intranasally challenged with a SARS-CoV-2 XBB.1.5 isolate (10^4.5^ TCID_50_ per animal). Naïve direct contact animals were separated for 24 hours (**Fig. 4a**). Following challenge, no mortality was observed across all vaccination regimens. However, significant body weight loss was detected in the mock-vaccinated group, with a clear difference compared to both the mRNA- or scVac^XBB^-group at 5 days post-challenge (dpc)—the peak of weight loss in the mock group (*p < 0.0001*). Moreover, scVac^XBB^-vaccinated animals gained more body weight than mRNA-vaccinated animals (*p = 0.0052*) (**Fig. 4b**). None of the direct contact animals experienced body weight loss after co-housing with challenged donors (**Fig. 4c**).

**Figure 4:**
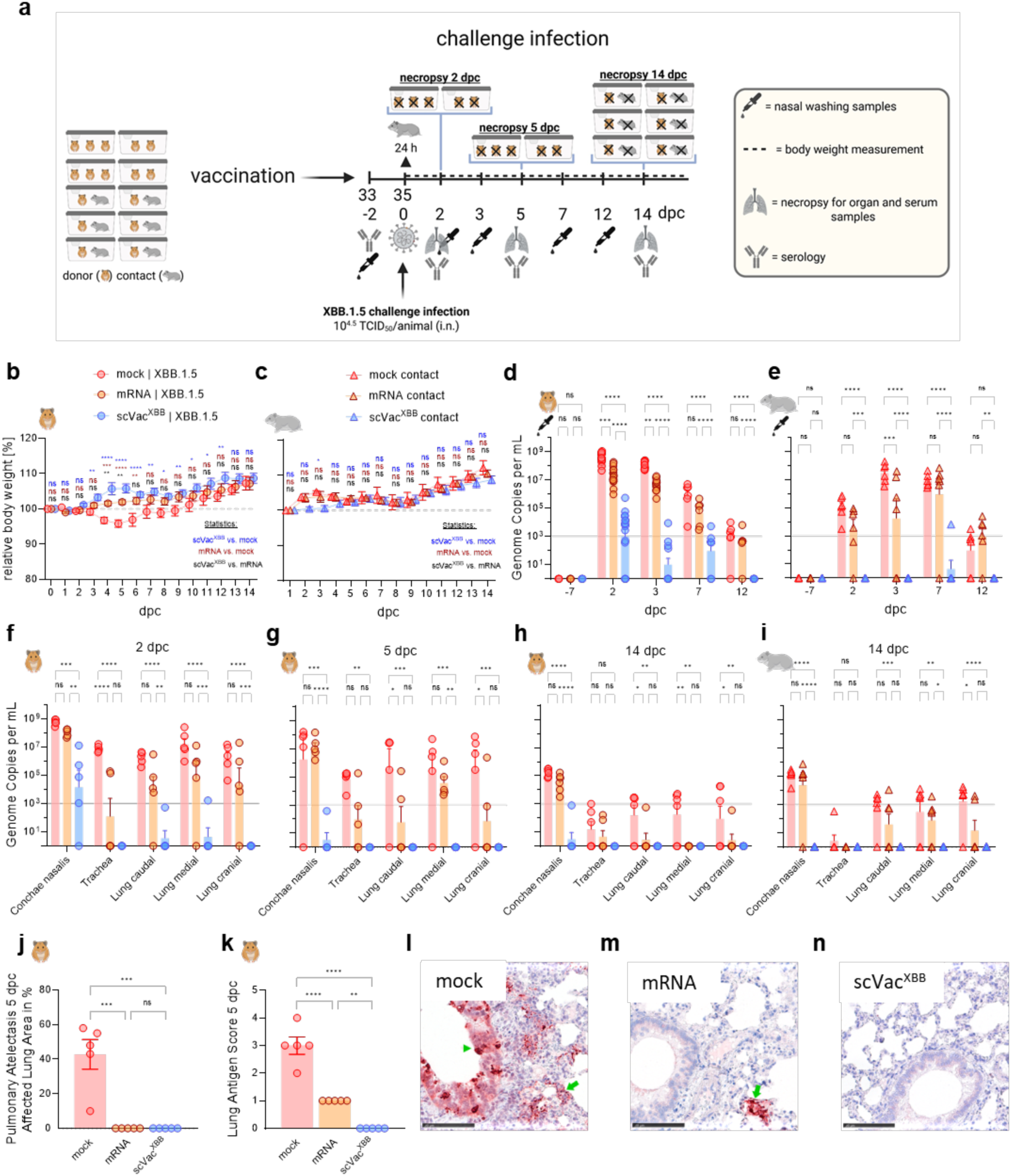
scVac^XBB^ protection from SARS-CoV-2 XBB.1.5 challenge infection. (a) Prime-boost vaccinated Syrian hamsters (mock, mRNA, scVac^XBB^; n = 16) were intranasally inoculated with SARS-CoV-2 XBB.1.5 at the time points as indicated. (b, c) Body weight as a function of time after challenge infection for (b) donor and (c) contact animals. (d-i) Quantification of SARS-CoV-2 genome copies in nasal washing samples before and after challenge infection in (d) donor and (e) contact animals and in lung tissue samples of donor hamsters at (f) 2, (g) 5, (h) and 14 dpc and of (i) contact animals at 14 dpc. (j) Pneumonia-related atelectasis, expressed as the percentage of affected area. (k) SARS-CoV-2 Nucleocapsid Antigen score in lung tissues. Representative images of SARS-CoV-2 nucleocapsid antigen in lung samples for (l) mock-, (m) mRNA-, and (n) scVac^XBB^-vaccinated, with indication of focal (mRNA) to multifocal (mock) presence of antigen in pneumocytes (green arrow) or bronchial epithelial cells (green arrowhead), scalebar is 100 µm. Statistical significance was calculated by ordinary one-way ANOVA with Tukey’s multiple comparison test. *\*p < 0.05, **p < 0.01, ***p < 0.001, ****p < 0.0001*, ns = non-significant. Besides individual values, group mean values are indicated by bars together with the standard error.

Virus shedding, measured in nasal wash samples, was significantly reduced in the scVac^XBB^ group compared to both mock (*p < 0.0001*) and mRNA-vaccinated animals (*p < 0.0001*) at all time points (**Fig. 4d**). Notably, none of the scVac^XBB^ direct contact animals showed evidence of productive infection (defined as positive detection at more than one time point), while all mock contacts and five out of six mRNA contact animals tested positive for challenge virus infection (**Fig. 4e**).

Only single animals in the scVac^XBB^-group displayed minimal viral genome presence at 2 and 5 dpc, with a complete absence by day 14, in contrast to mRNA and mock groups (**Fig. 4f-h**). Compared to mock-vaccinated animals, the challenge virus genome detections were significantly lower in the nasal conchae samples of scVac^XBB^-vaccinated animals (*p < 0.001* at 2 and 5 dpc, *p < 0.0001* at 14 dpc) and slightly reduced in mRNA-vaccinated animals (*p < 0.01* at 2, 5, and 14 dpc) (**Fig. 4f-h**). Evaluation of respiratory tissues at 14 dpc confirmed virus transmission to all mock contact animals and five out of six mRNA contacts, whereas none of the scVac^XBB^ contacts had detectable viral genomes in their lungs (**Fig. 4i**), indicating complete prevention of transmission.

Histopathological analysis revealed that all mock-vaccinated controls exhibited mild to moderate atelectasis and acute SARS-CoV-2-characteristic necrotizing bronchitis, vascular lesions, and marked alveolar edema with mixed cellular immune cell infiltration (**Fig. 4j, Supplementary Fig. 2a**). Intralesional, multifocal to coalescing viral antigen was detected in all hamsters (**Fig. 4k, l**).

In contrast, mRNA-vaccinated animals did not exhibit signs of pneumonia-induced atelectasis or acute bronchial or vascular lesions (**Fig. 4j, Supplementary Fig. 2b)**. However, viral antigen was present in a few foci in the alveolar epithelium of all animals, accompanied by signs of cell degeneration or necrosis with minimal heterophilic immune cell infiltration, consistent with acute infection (**Fig. 4k, m)**.

Similarly, no SARS-CoV-2-induced pulmonary atelectasis in the lungs of scVac^XBB^-vaccinated hamsters (**Fig 4j**). Furthermore, viral antigen was absent in these animals (**Fig. 4k, n**). No signs of acute infection, such as necrotizing bronchitis, intravascular rolling of immune cells, vasculitis, or infiltration by heterophilic granulocytes (**Supplementary Fig. 2c**). Both scVac^XBB^-and mRNA-vaccinated groups exhibited mild perivascular and peribronchial, predominantly lymphocytic immune cell infiltrates and interstitial immune cell expansion indicative of past immune stimulation (**Supplementary Fig. 2b, c**).

These findings demonstrate that scVac^XBB^ vaccination provides superior protection, effectively preventing both disease and transmission in a high-dose exposure model.

### Systemic and mucosal immunity are associated with transmission block

To examine the systemic immune response following challenge infection, serum samples collected at 2, 5, and 14 dpc were analyzed using a SARS-CoV-2 XBB.1.5 RBD-specific ELISA. scVac^XBB^-vaccinated animals exhibited a robust anamnestic (“boosting”) response as early as 2 dpc, significantly exceeding that of mRNA-vaccinated animals (*p < 0.0001*), while all samples from the mock-vaccinated group remained below the detection threshold (**Fig. 5a**). Complete blockade of virus transmission from scVac^XBB^-vaccinated animals to naïve direct contacts was confirmed by the absence of seroconversion in contact animals at 14 dpc. In contrast, only one mRNA contact animal remained seronegative, whereas all mock group contact animals seroconverted (**Fig. 5b**).

**Figure 5:**
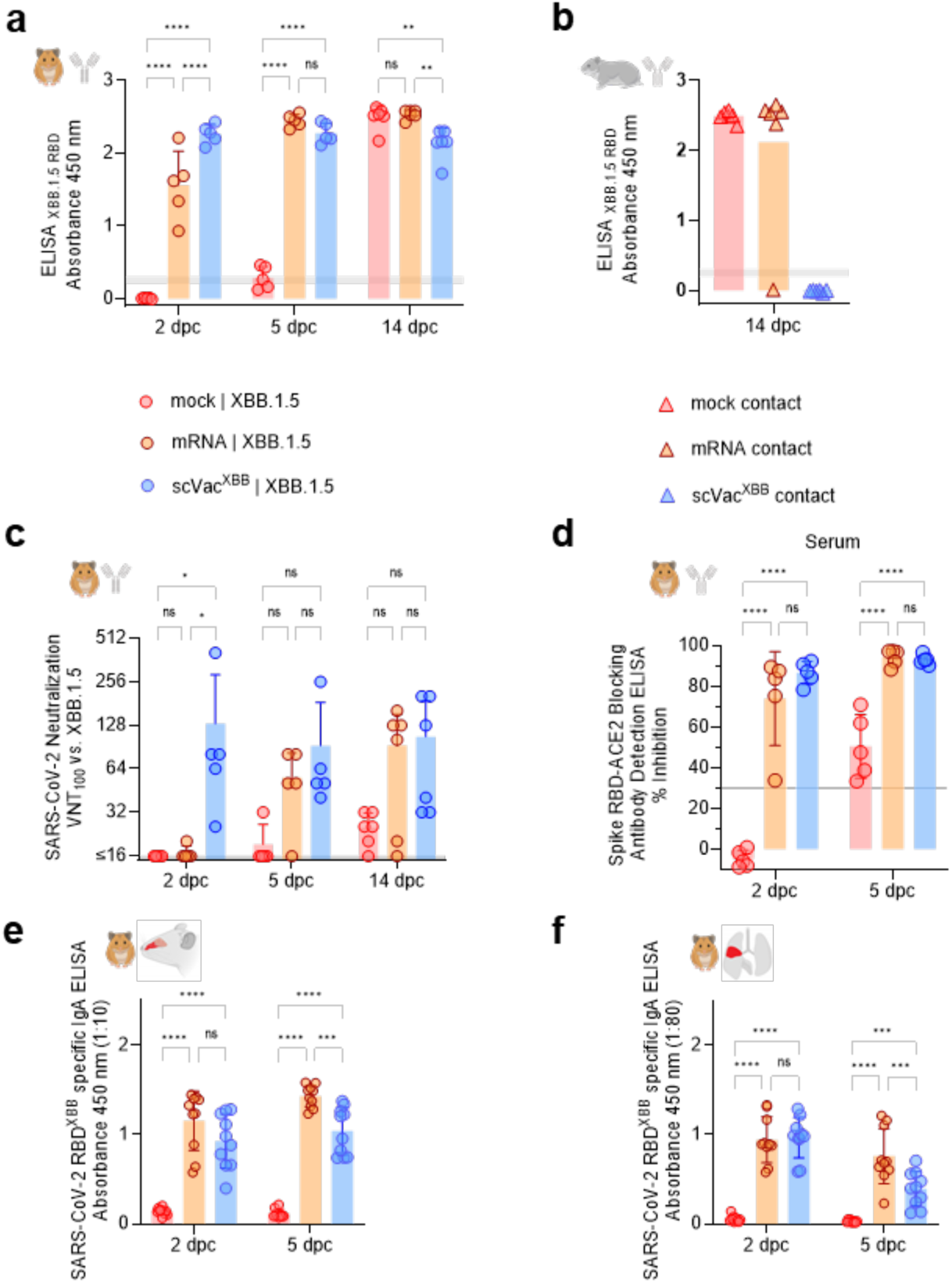
Mucosal and serum-related neutralizing immune response in vaccinated hamsters. (a, b) SARS-CoV-2 RBD-specific serum antibodies in (a) vaccinated and in (b) unvaccinated direct contact animals after challenge, measured by ELISA (samples were diluted 1:100). (c) Virus neutralization titer of serum at different time points after challenge to neutralize SARS-CoV-2 Omicron XBB.1.5 *in vitro*. (d) Detection of serum antibodies that were able to block RBD_XBB.1.5_-ACE2 binding by ELISA. (e, f) Detection of RBD_XBB.1.5_-binding IgA in (e) nasal and (f) lung tissue samples detected by ELISA. The full data set of dilution rows, ED_50_, and AUC values is shown in **Supplementary Fig. 4**. Individual values with bars indicating the mean with the respective standard error. Statistical significance was calculated by ordinary two-way ANOVA with Tukey’s multiple comparison test. *\*p < 0.05, **p < 0.01, ***p < 0.001, ****p < 0.0001*, ns = non-significant.

Neutralizing antibody titers in scVac^XBB^-vaccinated hamsters were detectable at 2 dpc (mean titer 131.4), while only one of five mRNA-vaccinated animals exhibitied neutralizing activity against SARS-CoV-2 XBB.1.5 (mean titer 20.16), and none was detected in the mock group (titer <16) (**Fig. 5c**). The neutralizing response in the mRNA-vaccinated group was delayed compared to the scVac^XBB^ group, while seroconversion in the majority of mock controls occurred by 14 dpc (**Fig. 5c**). At 2 dpc, both vaccines induced homologous neutralizing antibody responses, the bivalent mRNA (ancestral/BA.5) vaccine reacted against BA.5 (mean titer 47.9) and minimally against the ancestral strain (two animals with 20.16), while the scVac^XBB^ vaccine was reactive against XBB.1.5 (mean titer 131.4) (**Supplementary Fig. 3a**). By 5 and 14 dpc, the neutralizing responses were further elevated, with increasing divergence between groups. Here, a higher reactivity of sera from the bivalent mRNA (wild-type and BA.4/BA.5) vaccinated animals against D614G and BA.5 viruses was observed (**Supplementary Fig. 3b, c**).

High levels of serum antibodies that block ACE2-binding by attaching to the RBD were confirmed by ELISA in both scVac^XBB^ and mRNA-vaccinated animals 2 dpc. This was absent in the mock-vaccinated group, which began rising by 5 dpc (**Fig. 5d**). To assess mucosal immunity, RBD^XBB.1.5^-specific IgA was measured in nasal conchae (upper respiratory tract) and lung tissue (lower respiratory tract) samples. SARS-CoV-2-specific IgA was detectable in both nose and lung samples as early as 2 dpc with no significant difference between mRNA and scVac^XBB^-vaccinated animals **(Fig. 5e, f).** However, by 5 dpc, a stronger boost effect was observed for mRNA-vaccinated animals (**Fig. 5e, f**). There were no significant differences in ED_50_ values as well as for the area under the curve (AUC) values between mRNA- and scVac^XBB^-vaccinated animals in both nasal and lung samples at 2 dpc (**Supplementary Fig. 4a, b**) and 5 dpc (**Supplementary Fig. 4c, d**).

## Discussion

The development and characterization of the scVac^XBB^ SARS-CoV-2 vaccine candidate represent a significant advancement in the pursuit of safe and effective vaccination strategies against emerging variants of concern. Our findings demonstrate that the scVac-backbone is a promising and adaptable platform that allows efficient Spike protein updating. The Spike-independent genetic engineering strategy — targeted deletions of the E gene, ORF6, ORF7a, and truncation of ORF3a — confers a single-cycle infection profile rather than traditional virus attenuation. This feature enhances safety by minimizing residual infection and transmission risks typically associated with live-attenuated vaccines.

*In vitro* studies confirmed the single-cycle nature of scVac^XBB^ across multiple cell lines, demonstrating robust viral antigen expression without productive replication. While scVac^XBB^ was completely cleared in all cell lines tested here, we did observe low-level, prolonged virus detection in non-complementing Vero E6 cells under specific conditions. We interpret this as a likely consequence of nonphysiological circumstances, including the absence of a functional interferon response and potential replicon-like activity enabling cell-to-cell spread. Importantly, no productive replication was detected (i) *in vivo*, (ii) in *ex vivo* nasal epithelial cell cultures, or (iii) in other cell lines tested. These findings are in line with previous reports indicating that E gene deletion disrupts coronavirus assembly and release while preserving immunogenicity [12].

*In vivo* studies in K18-hACE2 transgenic mice supported these findings: scVac^XBB^-inoculated mice showed no signs of mortality, no weight loss, no detectable virus propagation, and no CNS pathology. This contrasts with SARS-CoV-2 XBB.1.5 infection and confirms the single-cycle nature of scVac vaccines [15]. Interestingly, no seroconversion was observed in this model, highlighting its limitations for assessing the immunogenicity of intranasal live vaccines. Despite this, these results underscore the excellent safety profile of single-cycle vaccines, positioning them as promising mucosal vaccine candidates for use in vulnerable populations, including infants, pregnant women, or immunocompromised individuals.

In the Syrian hamster model, a more natural model for SARS-CoV-2, the scVac^XBB^ vaccine reconfirmed the high safety profile and demonstrated a lack of transmission and strong immunogenicity. A potent humoral immune response, including neutralizing capacity, was evident post-boost. This aligns with known advantages of replicating vectors in inducing broad immune responses, particularly mucosal immunity, which is generally limited with current vaccine platforms [16]. In addition, the absence of scVac virus transmission events to co-housed naïve animals further emphasizes the biosafety of this vaccine candidate and thus a potential utility for immune-compromised individuals.

SARS-CoV-2, including the XBB.1.5 variant, has a remarkably high transmission rate in hamsters, even under airborne conditions [17]. In this regard, it is noteworthy that even in the stringent direct contact situation of our challenge study, all scVac^XBB^-vaccinated hamsters had developed a solid immunity against XBB.1.5 infection that was able to completely prevent any transmission to direct contacts. Viral load assessment showed a massively reduced viral genome copies in upper respiratory tissues and a complete absence of the virus in the lung. These findings highlight the superior protective capacity of scVac^XBB^ over a simultaneously tested current mRNA vaccine, which achieved only a partial reduction of viral replication and transmission [18]. Histopathology confirmed these results by proving the absence of viral antigen and acute lesions in scVac^XBB^-immunized hamsters, while oligofocal antigen-associated lesions remained observable in mRNA-vaccinated animals. The severity of the XBB1.5 challenge infection in non-vaccinated animals was evident through the presence of severe lung lesions associated with viral antigen.

Crucially, scVac^XBB^-induced immune responses correlated with complete prevention of virus transmission. This supports the notion that scVac can protect both the vaccinated individual and their contacts, likely mediated by strong mucosal immunity. Such features are essential for viral outbreak control [19] and are consistent with prior observations that LAVs can induce localized immunity capable of reducing viral shedding and secondary transmission [20]. However, the precise correlates of protection that facilitate these beneficial effects are not fully understood yet.

The successful integration of the epidemiologically relevant XBB.1.5 spike protein into the scVac backbone also underscores the platform’s adaptability to antigenic drift and shift, addressing concerns about immune evasion of newly emerging variants. This flexibility will remain beneficial for maintaining vaccine effectiveness against SARS-CoV-2 as this virus continues to evolve [21].

Future studies should evaluate the long-term durability of the induced immune response. Longitudinal studies will be critical to determine the persistence of protection and potential cross-protection against heterologous variants. Additionally, while Syrian hamsters and K18-hACE2 mice offer valuable preclinical insights, further validation in non-human primates and, ultimately, human trials will be necessary to confirm the translational potential of this vaccine platform.

## Methods

### Mouse holding and inoculation

An equal number of male and female K18-hACE2 transgenic mice (*Mus musculus*, B6.Cg-Tg[K18-ACE2]2Prlmn/J), aged 4 to 6 weeks, were procured from Charles River Laboratories. The mice were housed in groups of up to three within individually ventilated cages (IVCs) and were provided with unlimited access to food and water. Housing conditions were regulated at a temperature of 20-24 °C and humidity levels of 45-65%, with a 12-hour light/dark cycle that included a 30-minute dawn transition. Mice were intranasally inoculated with 25 µl of either SARS-CoV-2 Omicron XBB.1.5 (10^3.73^ TCID_50_/animal in 25 µl) (EPI_ISL_16640568) [22], with scVac^XBB^ (10^4.4^ TCID_50_/animal) or with non-infectious, 0.22 µm filtered cell culture supernatant. Under isoflurane-induced short-acting anaesthesia, the buccal cavity and nasal surface were swabbed with a moistened swab.

### Hamster handling and vaccination

Specific pathogen-free (SPF) Syrian hamsters (*Mesocricetus auratus*) were obtained from Janvier Labs (Le Genest-Saint-Isle, France). Equal numbers of male and female hamsters, aged 4-6 weeks, were used, totaling 66 animals. They were housed in groups of up to three per IVC, maintained at a temperature of 20-24 °C with 45-65% humidity. A 12-hour light/dark cycle with a 30-minute dawn and dusk transition was implemented. Standard rodent chow and water were provided ad libitum, along with enrichment materials such as straw and paper for nest building.

Following an acclimatization period, baseline measurements, including body weight and general health assessments, were recorded. The hamsters (n=16 per group) were vaccinated intranasally with scVac^XBB^ (10^4.89^ TCID_50_/animal) or mock (noninfectious, 0.22µm filtered cell culture supernatant) (70 µl were distributed equally in each nostril) or intramuscularly with a bivalent mRNA vaccine (Comirnaty, Original/Omicron BA.4-5 [23] [5 µg/hamster]). All animals were boosted under the same conditions 21 days following prime vaccination. The SARS-CoV-2 Omicron XBB.1.5 challenge virus (EPI_ISL_16640568) [22] was administered via intranasal inoculation with 10^4.49^ TCID_50_/animal in a 70 µl volume. To evaluate potential transmission of the live vaccine or the challenge virus, six of the vaccinated animals of each group were co-housed to serologically naïve direct contact hamsters in a 1:1 setup, after separating them for 24 hours following prime, boost, and challenge inoculation to exclude unspecific transmission of the inoculum.

Throughout 7 days post prime and boost vaccination, and 14 days following challenge infection, animals were monitored daily for clinical signs, weight changes, and overall health status. Samples were collected at predetermined intervals and included blood for serological analysis, nasal washing samples, and organ samples of the upper (conchae nasalis, trachea) and lower (caudal, medial, and cranial part of the left lung lobe) respiratory tract for viral load assessment, and lung samples (right lung lobe) for histopathological evaluation.

Nasal washing samples were taken at -2, 3, 7 dpv/dpb and at 2, 3, 7, and 12 dpc by applying 200 µL of PBS in each nostril and collecting the reflux under short isoflurane inhalation anesthesia. Blood samples were either taken at end timepoints (2, 5, and 14 dpc) or by puncturing the V. saphena at -2, 19, and 33 dpv for serological evaluation.

### Cell lines

African green monkey kidney cells (Vero E6) were kindly provided by V. Thiel, Bern, Switzerland, or obtained from the Collection of Cell Lines in Veterinary Medicine CCLV-RIE 0929. Adenocarcinomic human alveolar basal epithelial cells expressing ACE2 and TMPRSS2 (A549-AT) were obtained from NIBSC (A549-ACE-2 Clone 8-TMPRSS2; product number 101006). HEK293T cells were kindly provided by D. D. Pinschewer, University of Basel, Switzerland, and CaCo2 cells were provided by G. Kochs, University of Freiburg, Germany.

All cells were maintained in DMEM high glucose with 10% FBS + 1% Penicillin / Streptomycin for general propagation or with 2% FBS + 1% Penicillin / Streptomycin for viral infection experiments. During the initial viral rescue, HEK293T indE were maintained with the JAK-I inhibitor Pyridone 6 (CAS 457081-03-7) at a final concentration of 2 µM as well as the NFκB inhibitor QNZ (CAS 545380-34-5) at 20 nM.

### Cell line generation

HEK293T-indE cells (HEK293T-E Tet::E-IRES-ORF6) are a derivative of HEK293T-E cells (stable expression of the SARS-CoV-2 E gene under CMV promoter) with a second-generation lentiviral vector generated with the pCW57-E-IRES-ORF6 (Addgene plasmid #80921) as a transfer vector. The vector codes for SARS-COV-2 E and ORF6 under a Tetracycline inducible promoter. Cells were analyzed by RT-qPCR for E and ORF6 induction following doxycycline treatment [13].

Vero-E2T cells were generated by transfecting Vero E6 cells with 2 µg of an equimolar plasmid mixture containing the SARS-COV-2 E/ORF6/ORF7a/ORF8 genes in individual plasmids all under the CMV promoter in a pcDNA3.1 background. Human TMPRSS2 expression in Vero-E2T and Vero E6 cells (Vero E6-TMPRSS2) was achieved by infecting the cells with a 2nd generation lentiviral vector pLEX307-TMPRSS2-blast (Addgene plasmid #158458) as a transfer vector. For details on transgene expression, see [13].

### Viral genome reconstitution procedures

Virus recovery was achieved as outlined in [14]. In brief, PCR fragments (fr A, B, C, D1, and D2) encompassing the entire SARS-CoV-2 genome were amplified using the high-fidelity proofreading enzyme Q5^®^ High-Fidelity DNA Polymerase (NEB, M0491L) in a 25 µL reaction volume with the respective primers. Fragment A includes the heterologous CMV promoter upstream of the 5’ UTR, while fragment D2 contains the poly(A) tail, HDV ribozyme, and SV40 termination signal downstream of the 3’ UTR (Fig. 1a).

Cycling conditions were used as recommended by the manufacturer. Fragments were obtained using the following primer combinations: frA: CMV for + frA-frB rev; frB: frB-frA for + frB-frC rev; frC: frC-frB for + frC-frD rev; frD1: frD-frC for + S rev and frD2: frD2 for + SV40 rev. DNA oligonucleotide sequences used are listed in Supplementary Table 1.

12-30 reactions were pooled and purified by PCR column purification using the QIAquick PCR purification kit (Qiagen, 28104). DNA concentration was measured by Nanodrop 1000 (Thermo Fisher) or Quantus (Promega, QuantiFluor® ONE dsDNA System, E4871). DNA was further purified by ethanol precipitation, and the final concentration was adjusted to 1 µg / µL in nuclease-free water.

Equimolar ratios of frA, frB, frC, frD1, and frD2 and all-in-E plasmid were transfected into HEK293T-indE using jetPRIME® (Polyplus, cat. 101000001) as recommended by the manufacturer. 4-24h post-transfection, the medium was changed to DMEM with 2% FBS with the addition of JAK-I inhibitor Pyridone 6 (CAS 457081-03-7) to a final concentration of 2 µM, as well as the NFκB inhibitor QNZ (CAS 545380-34-5), and Vero-E2T cells were added for co-incubation. Every 3-4 days, the medium was exchanged. Screen for virus progeny production was done with SARS-CoV-2 antigen rapid tests (Roche, 9901-NCOV-01G, and Clungene®, GTIN: 6950921302797) or by cytopathic effect (CPE) in Vero-E2T cells and confirmed by RT-qPCR and FFU (focus forming unit) quantification.

### Virus propagation for storage

For wild-type controls, clinical isolates XBB.1.5 (isolated from nasopharyngeal aspirates of human donors who had given their informed consent or received from Bart Haagmans, Rotterdam [EPI_ISL_16640568]), rCoV2 (recombinant Wuhan-1-type virus produced by genome reconstitution [13,14]) or rCoV2^XBB^ (recombinant Wuhan-1-type virus with Omicron XBB.1.5 Spike generated with CLEVER), were propagated in Vero E6 cells until CPE was observed. For deletion mutants, viral particles produced by HEK293T-indE were further amplified in Vero-E2T cells. Viral propagation was observed and monitored by CPE and antigen rapid tests and confirmed by RT-qPCR and FFA.

Final viral stocks were harvested, filtered by 0.2 µm filters to remove cells, and frozen in small aliquots. For each viral stock, the viral titer was determined by RT-qPCR and focus forming assay (FFA).

### RNA extraction for virus quantification and sequencing of viral stocks

Viral RNA was extracted using the automated Promega Maxwell RSC system (Promega, AS4500) using either the Maxwell® RSC Viral Total Nucleic Acid Purification Kit (Promega, AS1330) or the Maxwell® RSC miRNA from the Tissue and Plasma or Serum Kit (Promega, AS1680).

### Dideoxy-sequencing

The region of interest was amplified using SuperScript™ IV One-Step RT-PCR System (Thermo Fisher, 12594100) with either ‘frD2 for’ or ‘26847 for’ and ‘29046 N rev’ (Supplementary Table 1). The integrity of the PCR product was checked on agarose gel and subsequently sent for Sanger sequencing to evaluate genome regions affected by deletions/mutations (Microsynth, Switzerland).

### Next-generation sequencing (NGS)

Viral RNA was converted to cDNA using a cDNA Synthesis kit (biotechrabbit). cDNA was NGS sequenced using EasySeq SARS-CoV-2 WGS Library Prep Kit (NimaGen, SKU: RC-COV096) on an Illumina NextSeq 2000 system with a P1 flow cell (300 cycles). All NGS sequencing and raw data analysis was done by Seq-IT GmbH & Co. KG.

### RT-qPCR quantification of viral and intracellular RNA

For the detection of SARS-CoV-2 RNA, a primer and TaqMan probe set for ORF-1b (Supplementary Table 1) were used as described [24]. The RT-qPCR Luna® Universal Probe One-Step RT-qPCR Kit (E3006E) was used according to the manufactureŕs protocol. In brief, Master Mix was set up: for one reaction, 1 µL of each primer, 0.5 µL Probe, 10 µL of Luna Universal Probe One-Step Reaction Mix (2X), 1 µL of Luna WarmStart RT Enzyme Mix (20X) were mixed and brought to15 µL with nuclease-free water. 15 µL of Master Mix was mixed with 5 µL RNA and amplified on an ABI7500 fast cycler (ThermoFisher) using the following cycling conditions: 10 min 55 °C, 1min 95°C denaturation, followed by 45 cycles for 10 seconds at 95°C and 30 seconds at 60°C.

### In vitro passaging for in vitro safety experiments

For viral passaging experiments, Vero E6, CaCo2, or A549 ACE2/TMPRSS2 cells were infected in duplicates with an MOI of 0.01 (based on FFU) for 3-4h with *wild-type* virus or deletion candidate. The cells were washed, and 2% DMEM medium was added. Every fourth day, supernatant (SN) was passaged on freshly seeded cells (25% confluency). SN was always diluted 1:100. All respective passages were extracted, and RT-qPCR was performed to quantify residual viral RNA. All conditions were treated equally.

### Immunocytochemistry

For detection of infectious vaccine viral particles (FFA) and protein expression analysis, Vero E6-TMPRSS2 cells grown on coverslips in 24-well plates were infected with virus variants in 500 µL DMEM medium supplemented with 2% FCS and 1% Penicillin / Streptomycin and incubated overnight. Cells were fixed with 4% PFA in PBS for 10 min at room temperature, washed, and subsequently stained. Cells were blocked with 10% Normal Donkey Serum (Jackson ImmunoResearch, 017-000-121) and 0.1% Triton X-100 at room temperature for 60 min followed by incubation with primary antibodies for 60 min at room temperature or overnight at 4°C in 1% Normal Donkey Serum, 1% BSA and 0.3% Triton X-100 in PBS. Cells were washed three times for 10 min with 0.1% BSA / PBS and incubated with fluorophore-coupled secondary antibodies for 60 min at room temperature in 1% Normal Donkey Serum, 1% BSA, and 0.3% Triton X-100 in PBS. Cells were washed once with 0.1% BSA / PBS and washed three times with PBS before mounting on microscope slides using Fluoromount-G (SouthernBiotech, 0100-01). Phalloidin-iFluor488 was co-applied with secondary antibodies to label F-actin (Abcam, ab176753). Hoechst 33342 dye (Merck, B2261) was co-applied during washing at a final concentration of 0.5 µg / mL for nuclear staining.

Images for FFU quantification were acquired on a bright-field microscope (Nikon Ti2 equipped with a Photometrics 95B camera, Nikon NIS AR software), using a 20x Plan-Apochromat objective (numerical aperture 0.75) and were then processed in Fiji and Omero. For quantification of infected foci, images were analyzed with QuPath [25]. Images for protein expression were acquired on an inverted spinning-disk confocal microscope (Nikon Ti2 equipped with a Photometrics Kinetix 25mm back-illuminated sCMOS, Nikon NIS AR software), using a 40x Plan-Apochromat objective (numerical aperture 0.95 and 1.45 respectively) and were then processed in Fiji and Omero.

### Biochemical procedures

For validation of the vaccine candidate, Vero E6-TMPRSS2 cells were infected with virus variants at an MOI of 0.1. 24h after infection, cells were washed twice with PBS before lysis in cold 140 mM NaCl, 50 mM Tris-HCL, 1% Triton-X100, 0,1% SDS, 0,1% sodium deoxycholate supplemented with protease and phosphatase inhibitors (ThermoFisher, 1861281). Lysates were centrifuged for 10 min, 16’000g at 4°C, and supernatants were analyzed by Immunoblot. Signals were acquired using an image analyzer (Odyssey CLx, Licor).

### Antibodies

The following antibodies were used for immunocytochemistry and immunoblotting: mouse monoclonal anti-β-actin (Cell Signaling Technology; 3700; RRID: AB_2242334; LOT# 20), rabbit polyclonal anti-SARS-CoV-2 nsp2 (GeneTex; GTX135717; RRID: AB_2909866; LOT# B318853), mouse monoclonal anti-SARS-CoV-2 Nucleocapsid protein (4F3C4, gift from S. Reiche [26], sheep polyclonal anti-SARS-CoV-2 ORF3a [27], rat monoclonal anti-SARS-CoV-2 ORF6 (8B10, gift from Y. Miyamoto [28]), mouse monoclonal anti-SARS-CoV-2 ORF7a (3C9; GeneTex; GTX632602; RRID: AB_2888320; LOT# 42219), rabbit polyclonal anti-SARS-CoV-2 ORF8 (Novus Biologicals; NBP3-07972; LOT# 25966-2102), mouse monoclonal anti-SARS-CoV-2 Spike protein (4B5C1, gift from S. Reiche).

Fluorophore-conjugated secondary antibodies were from Jackson ImmunoResearch (Cy3 donkey anti-mouse #715-165-151, Cy3 donkey anti-rabbit #711-165-152, Cy5 donkey anti-rat #712-175-153, Cy5 donkey anti-mouse #715-175-511, Cy5 donkey anti-sheep #713-175-147), Li-Cor (IRDye 680RD donkey anti-mouse #926-68072, IRDye 680RD goat anti-rabbit #926-68071, IRDye 680RD goat anti-rat #926-68076) and Invitrogen (Alexa Fluor 680 donkey anti-sheep #A21102).

### Genome copy quantification by RT-qPCR (animal experiments)

A total of 100 µl of nasal washing or organ samples were utilized for the extraction of nucleic acids, employing the NucleoMag Vet kit (Macherey-Nagel). To quantify SARS-CoV-2-specific nucleic acids, the RdRp locus was targeted [29]. To confirm successful RNA extraction, the housekeeping gene β-actin [30] was used (for primers see Supplementary Table 1). The RT–qPCR reactions were prepared using the qScript XLT One-Step RT–qPCR ToughMix (QuantaBio) in a total volume of 12.5 μl, including 1 μl of the FAM mix and 2.5 μl of extracted RNA. The amplification was performed on a BioRad real-time CFX96 detection system (Bio-Rad) under the following cycling conditions: 50°C for 10 minutes, denaturation at 95°C for 1 minute, followed by 42 cycles of 10 seconds at 95°C, 10 seconds at 60°C, and 20 seconds at 68°C. Fluorescence was measured during the annealing phase.

### Virus titration

The virus’s tissue culture infectious dose (TCID_50_/mL) was determined using standard endpoint dilution assays on Vero E6 cells. In brief, the cells were plated in a 96-well plate one day before titration, reaching 80–90% confluency. Subsequently, they were inoculated with 100 µL of a 10-fold serial dilution of the sample. Each dilution was rigorously tested in 16 technical replicates. The cells were then incubated at 37°C in a humidified incubator with 5% CO^2^ for 72 hours. Virus titers were calculated using the midSIN calculation [31].

### RBD-Specific Enzyme-Linked Immunosorbent Assay (RBD-ELISA)

The RBD-ELISA was performed following the protocol described in [32]. Binding plates (Greiner, Kremsmünster, Austria) were coated overnight at 4 °C with 100 ng/well of SARS-CoV-2 RBD antigen in 0.1 M carbonate buffer. Afterward, the plates were blocked at 37 °C for 1 hour using 5% skim milk in PBS. Serum samples, diluted 1:100 in Tris-buffered saline with Tween 20 (TBST), were incubated on the coated wells for 1 hour at 37 °C. A multi-species conjugate (SBVMILK; derived from the ID Screen® Schmallenberg Virus Milk Indirect ELISA; IDvet) diluted 1:80 was subsequently added. Reactivity was visualized with the addition of tetramethylbenzidine (TMB) substrate (IDEXX), using a Tecan plate reader (Tecan Group Ltd, Männedorf, Switzerland) with 450 nm wavelength detection. Plates were washed three times with TBST between each step. Each assay included two replicates of defined positive (PC) and negative (NC) control sera to ensure consistency and reliability of the results.

### IgA determination by ELISA

SARS-CoV-2 XBB.1.5 Spike protein specific IgA was detected in serum and supernatants of homogenates from hamster conchae and lung adapting a protocol described in [10]. ELISA plates (96-well, flat-bottom; Nunc MaxiSorp) were coated with 100 µl of 1.5 µg ml−1 SARS-CoV-2 XBB.1.5/Omicron Spike Trimer Protein (Arco Biosystems) in PBS overnight at 4 °C. The following day, plates were washed three times with PBS supplemented with 0.05% Tween 20 (PBS-T) and incubated with 5% milk in PBS (blocking buffer) for 1 h at RT to block unspecific binding. Organ homogenates were centrifuged at 4,000 × g for 5 min. Samples were serially diluted 3-fold in blocking buffer starting at at 1:10 dilution for supernatants and 1:100 dilution for serum, before adding 50 µl of the diluted samples to the plates. Samples were incubated in the plates for 2 h at r.t., then washed three times before adding 50 µl of 1:50-diluted biotinylated anti-hamster IgA detection antibody (Brookwood Biomedical). Following a 2 h incubation at r.t., plates were washed three times and 50 µl of High-Sensitivity NeutrAvidin HRP conjugate was added for 30 min at r.t. The plates were washed three times and 50 µl of 1-Step Ultra TMB ELISA substrate solution (Thermo Fisher) was added. After 5 min, the reaction was stopped by adding an equal volume of 2 M sulfuric acid. The plates were read for absorbance at 450 nm and 570 nm on a Tecan Infinite M200 Pro microplate reader. Extinction at 570 nm was subtracted as background. The effective dilution to reach 50% of the maximal extinction (ED_50_) for each sample was determined using a four-parameter nonlinear regression curve fit in GraphPad Prism v.9 as well as the area under the curve (AUC) which reflects the cumulative antibody binding signal across the dilution range.

### Virus Neutralization Test (VNT)

The neutralizing capacity of serum samples against live SARS-CoV-2 XBB.1.5 was evaluated in vitro using Vero E6 cells (Vero C1008) cultured in 96-well plates (Greiner, Kremsmünster, Austria). Serum samples were diluted two-fold in triplicate and mixed with an equal volume of SARS-CoV-2, corresponding to 100 TCID_50_ per well. Used SARS-CoV-2 variants: An ancestral B.1 isolate BavPat1 (SARS-CoV-2 D614G mutation (GISAID number: EPI_ISL_406862)), an Omicron BA.5 VOC (EPI_ISL_12268493), and the Omicron XBB.1.5 VOC (EPI_ISL_16640568). Following a 1-hour incubation at 37 °C, 100 µL of trypsinized cells (harvested from a confluent 75 cm^2^ flask and suspended in 50 mL DMEM supplemented with 2% penicillin/streptomycin) were added to each well. The plates were incubated at 37 °C for three days, after which the presence of cytopathic effect (CPE) was assessed using a standard optical transmission microscope.

The neutralization titer (NT) was determined from three independent dilutions that fully inhibited CPE. The titer was calculated with the Kerber formula: −log2=𝑎/𝑏+𝑐, where 𝑎 represents the number of wells without detectable virus replication, 𝑏 is the number of wells tested per serum dilution, and 𝑐 is the −log2 value of the serum pre-dilution. The final titer, expressed as the −log2 of the highest serum dilution that completely prevented CPE in Vero E6 cells infected with 100 TCID_50_, is reported as the virus neutralization titer 100 (VNT_100_).

### Histopathology and immunohistochemistry of lungs

For histopathology, the left lung lobe was processed as described previously [33]. Briefly, the left lung lobe was carefully removed, immersion-fixed in 10% neutral buffered formalin, embedded in paraffin, and 2-3-μm sections were stained with hematoxylin and eosin (HE). Consecutive sections were processed for immunohistochemistry (IHC) according to standardized procedures of the avidin-biotin-peroxidase complex (ABC)-method. A primary antibody against the SARS-CoV Nucleocapsid protein was applied overnight at 4 °C (Rockland, 200-401-A50, 1:3000, RRID: AB_828403), and the secondary antibody, a biotinylated goat anti-rabbit antibody, was applied for 30 min at RT (Vector Laboratories, Burlingame, CA, USA, 1:200). As a negative control, consecutive sections were labelled with an irrelevant antibody (Glial Fibrillary Acidic Protein (GFAP), Abcam, ab16997, 1:600, RRID: AB_443592). In each run, an archived control slide from a SARS-CoV-2-infected Syrian hamster was included. All slides were scanned with a Hamamatsu S60 scanner and analyzed by a trained pathologist (T.B.) and reviewed by a certified pathologist (A.B.) in a blinded manner using NDPview.2 plus software (Version 2.8.24, Hamamatsu Photonics, K.K. Japan). The lung tissue was evaluated using a 500 × 500 μm grid, and the extent of pneumonia-associated consolidation was recorded as a percentage of the affected lung fields. Furthermore, the lung was examined for the presence of SARS-CoV-2-characteristic lesions described for hamsters. These included intra-alveolar, interstitial, peribronchial and perivascular inflammatory infiltrates, alveolar edema, necrosis of the bronchial epithelium, diffuse alveolar damage, vasculitis, activation of endothelium with immune cell rolling, as well as bronchial epithelial and pneumocyte type 2 hyperplasia. Following IHC, the distribution of virus antigen was graded on an ordinal scale. The scores were as follows: 0 = no antigen, 1 = focal, affected cells/tissue <5% or up to 3 foci per tissue; 2 = multifocal, 6%–40% affected; 3 = coalescing, 41%–80% affected; 4 = diffuse, >80% affected. The target cell was identified based on its morphological characteristics.

## Data Availability

All data generated or analyzed during this study are included in this published article and its supplementary information files.

## Ethical and Biosafety Statement

All procedures complied with institutional and national guidelines for the care and use of animals in research. The study protocol was approved by the relevant ethical review board (LVL MV TSD/7221.3-1-041/20). All work involving SARS-CoV-2 and its descendants was conducted in a biosafety level 3 (BSL-3) facility, adhering to established containment procedures.

## Acknowledgements

The research was supported by a grant of the German Research Foundation (DFG) (project number 453012513 / BE-5187/7-1), through an Innosuisse grant (IP-LS 119.403), and by RocketVax Inc. The authors thank the DBM Microscopy Core Facility for support with image acquisition and analysis. We thank M. Grawe for excellent technical assistance with analyzing the samples from the in vivo experiments and S. Schuparis for excellent technical assistance with the pathological analyses. We are also grateful to F. Klipp, S. Kiepert, C. Lipinski, D. Fiedler, and H. Manthei for their excellent care and dedication throughout the *in vivo* experiments and to Martin Daeumer and Alex Thielen (Seq-It GmbH, Kaiserslautern, GER) for expert NGS support. Vero E6 cells were kindly provided by Volker Thiel, Caco2 cells by Georg Kochs. Graphical symbols in Figures 2-5 were created with BioRender.com.

## Author contributions

Conceptualization: J.S., T.K., Do.H., F.O., M.B., Da.H.

Methodology: J.S., B.C., Do.H., F.O., Da.H.

Investigation: J.S., N.J.H., T.B., L.U., J.K., E.T.K., Do.H., F.O., Da.H.

Formal analysis: J.S., T.B., A.B., F.O., Da.H.

Data curation: J.S., N.J.H., F.O. Visualization: J.S., Da.H. Writing – original draft: J.S.

Writing – review & editing: J.S., N.J.H, T.B., A.B., J.K., B.C., E.T.K., T.K., Do.H., F.O., M.B., Da.H.

Funding acquisition: T.K., Do.H., F.O., M.B., Da.H.

Project administration: J.S., Do.H., F.O., M.B., Da.H.

Supervision: A.B., Do.H., F.O., M.B., Da.H.

All authors read and approved the final manuscript.

