## Supplementary Figures for "One and Done: A safe, adaptable single-cycle SARS-CoV-2 vaccine platform blocks XBB.1.5 infection and transmission"

**This PDF file includes:**

*Supplementary Figures 1 to 4:*

Supplementary Figure 1: scVac$^{XBB}$ viral titer quantification

Supplementary Figure 2: Histopathology of hamster lungs at 5 dpc

Supplementary Figure 3: Live virus neutralization tests

Supplementary Figure 4: IgA detection in nasal conchae and lung tissue samples

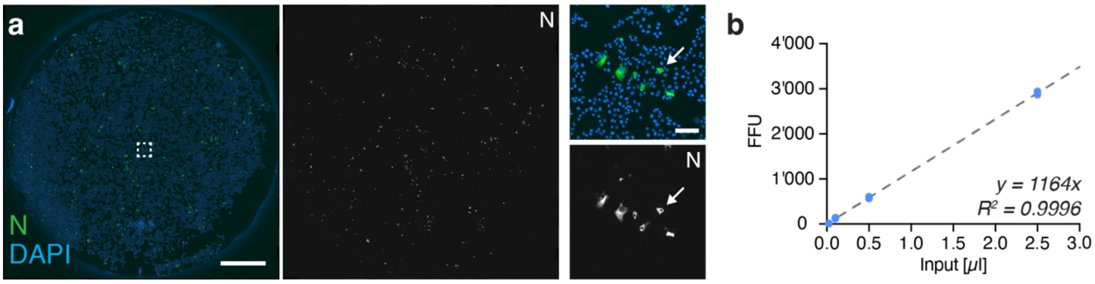


**Supplementary Figure 1: scVac$^{XBB}$ viral titer quantification**

(a) Representative example of infected cells used to quantify viral titers of scVac$^{XBB}$ on coverslips in a 24-well culture dish in Vero E6-TMPRSS2 cells, detection of the Nucleocapsid protein is shown in green, nuclei are stained with Hoechst (blue). Left: overview images, right: region of interest images showing individual infected cells as indicated. (b) Titration of scVac$^{XBB}$ and quantification by FFA (n = 2). Linear fit and correlation indicated for viral titer used to inoculate Syrian hamsters.

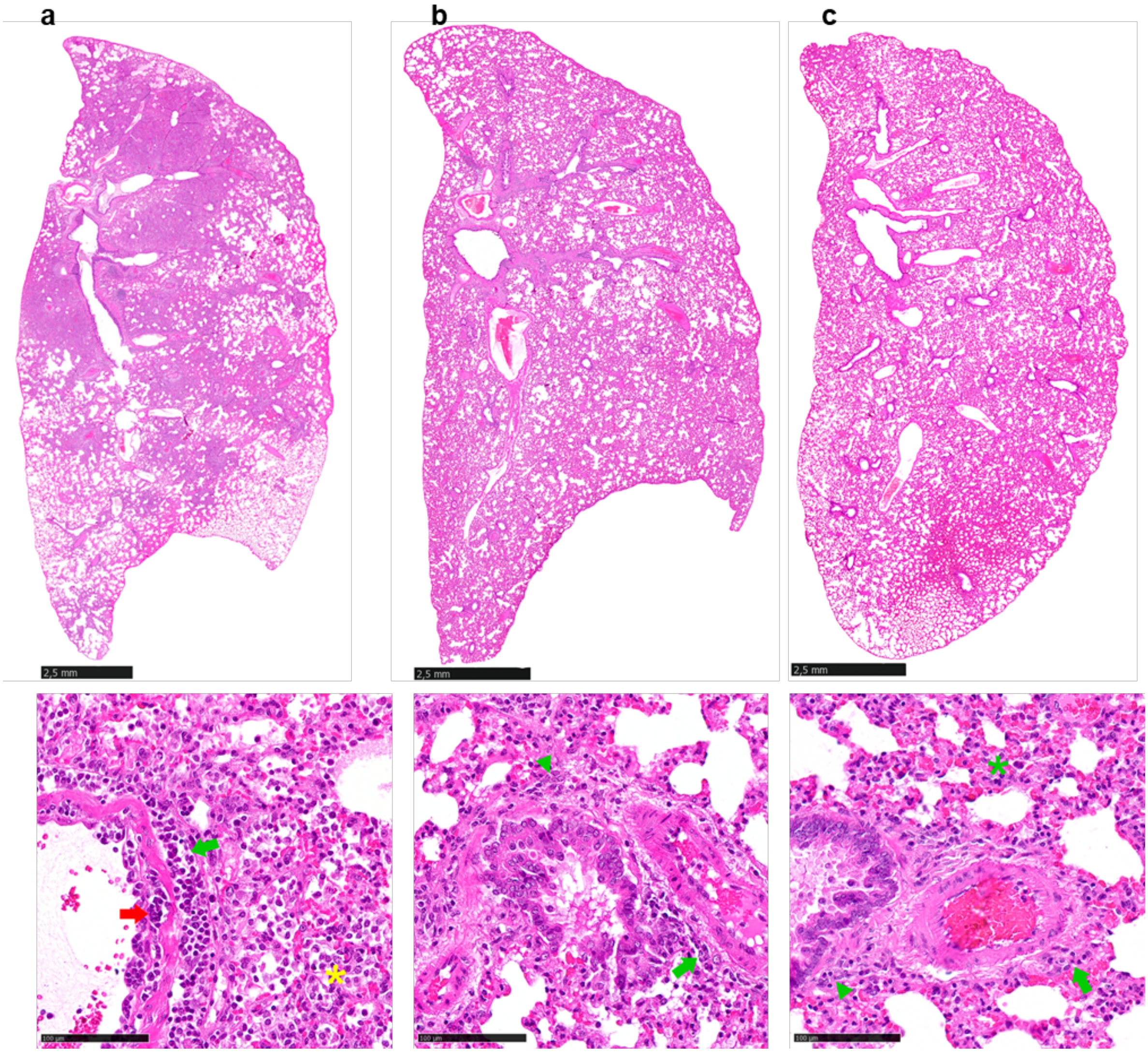


**Supplementary Figure 2: Histopathology of hamster lungs at 5 dpc.**

Representative hematoxylin-eosin-stained lung sections of the left lung (upper panels, bar 2.5 mm) and detail images (lower panels, bar 100 µm). (a) Non-vaccinated mock hamsters showed moderate atelectasis, along with marked perivascular inflammatory infiltrates (green arrow), vasculitis (red arrow), alveolar edema and mixed alveolar immune cell infiltration (yellow asterisk), while (b) mRNA- and (c) scVac$^{XBB}$-vaccinated hamsters showed no atelectasis and only mild perivascular (green arrow) and peribronchial (green arrowhead), mainly lymphocytic inflammatory infiltrates and a slight expansion of the interstitium by these immune cells (green asterisk).

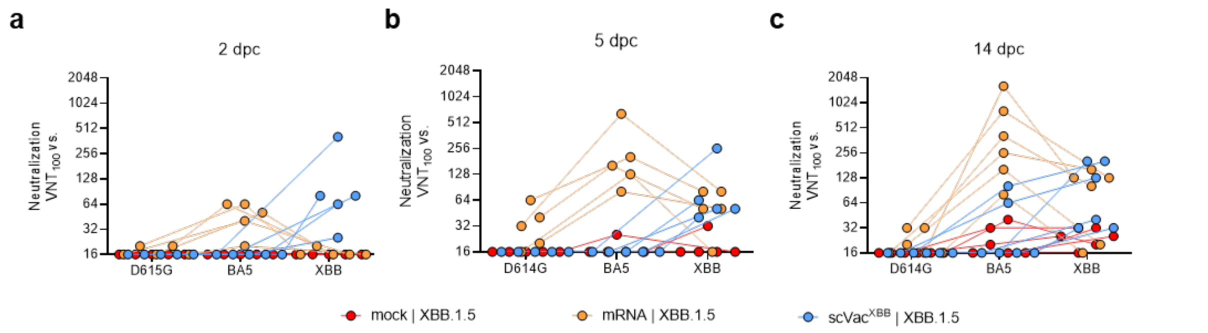


**Supplementary Figure 3: Live virus neutralization tests**

Serum samples obtained (a) 2, (b) 5, and (c) 14 days post challenge were tested against ancestral (D614G [BavPat1]) and Omicron BA.5 or XBB.1.5 SARS-CoV-2.

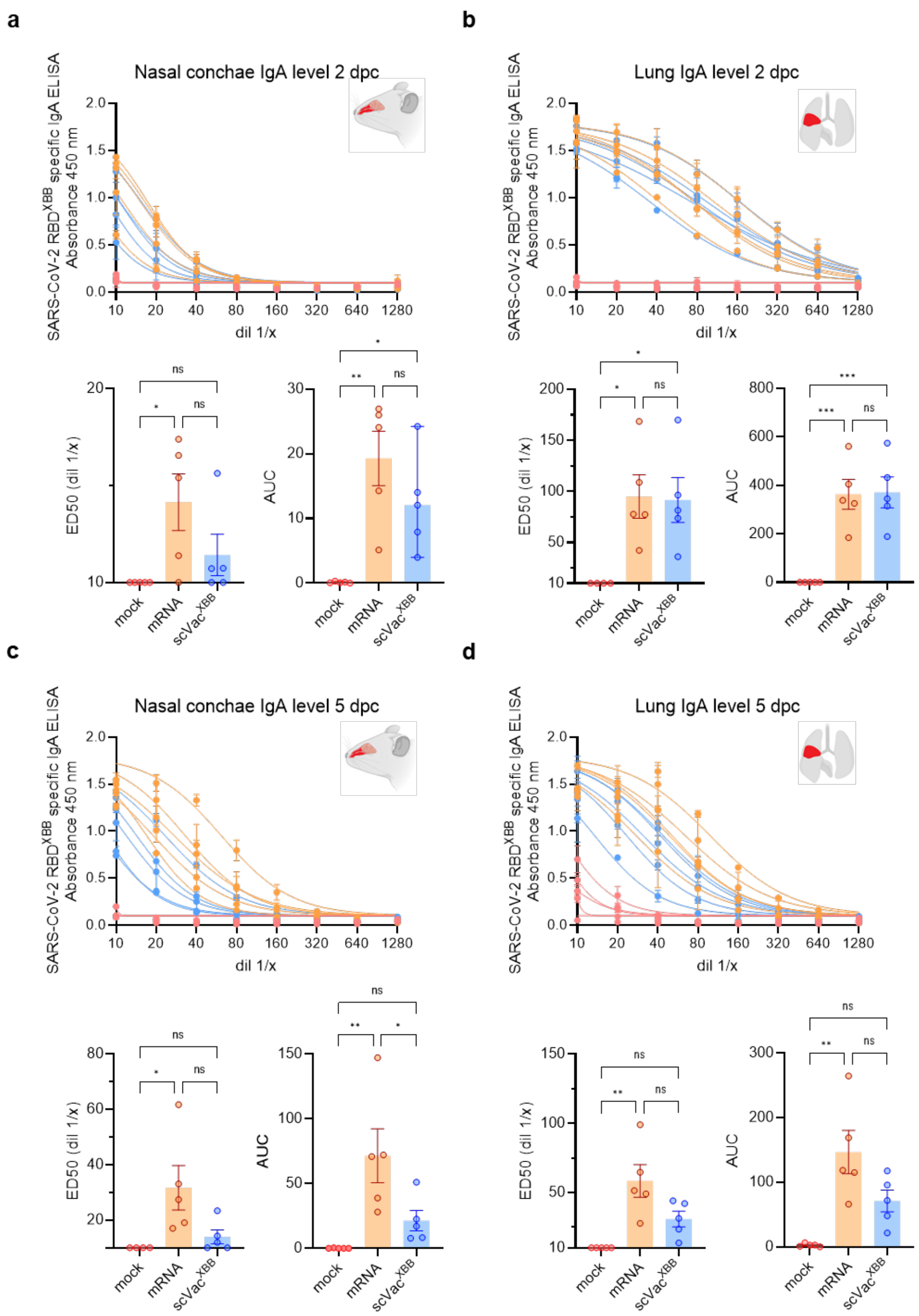


**Supplementary Figure 4: IgA detection in nasal conchae and lung tissue samples.** Dilution rows of (a) 2 dpc conchae and (b) lung sample homogenates and (c) 5 dpc conchae and (d) lung sample homogenates were measured by SARS-CoV-2 XBB.1.5 spike-coated ELISA specific for hamster IgA. Each dilution was measured in two technical replicates, and the mean was plotted, with each line representing one animal. The effective dilution to reach 50 % ($ED_{50}$) of the maximal dilution was calculated for each sample. It quantifies the antibody titer, i.e., the dilution at which antibody binding activity is still at 50% of the peak response. Furthermore, the area under the curve (AUC) was calculated, a relative measure of the total antibody response based on OD values across different dilutions. Statistical significance was calculated by ordinary one-way ANOVA with Tukey´s multiple comparison test. *$p < 0.05$, **$p < 0.01$, ***$p < 0.001$, ****$p < 0.0001$, ns = non-significant.
